# A method for analyzing physiological data with multiple non-independent observations

**DOI:** 10.1101/2025.06.25.661570

**Authors:** Erin C. McKiernan, Jorge Humberto Arce Rincón, Araceli Torres Pérez, Marco A. Herrera Valdez

## Abstract

Physiological studies often involve recording multiple observations (e.g., repeated muscle contractions or cardiac cycles) from the same subject. To compare groups of subjects, observations from one subject are then sometimes pooled with observations from other subjects in the same group, and analyses are performed on these pooled data. This approach presents a number of potential problems, including over-or under-representing certain subjects in the sample and non-independence of measurements. There are statistical methods available to deal with repeated measures, but many make assumptions about the distribution and completeness of data. Instead, we developed a method to deal with multiple observations from physiological data that ensures that each subject is represented only once in the analysis (i.e. it avoids pseudoreplication), and makes no assumptions about the distribution of the data. We demonstrate this method using two different physiological datasets: (1) muscle recordings taken from *Drosophila melanogaster* larvae during fictive crawling, and (2) electrocardiogram recordings taken from human volunteers before, during, and after exercise. Our results show the broad applicability and validity of this method.

## Introduction

The study of physiological processes, like motor control or heart rate variability, often requires taking multiple observations from the same subject. For example, studying fictive crawling in *Drosophila melanogaster* larvae involves intracellular or extracellular recording of multiple muscle contraction cycles [1, 2]. This spontaneous activity can last from seconds to minutes, representing tens of cycles recorded from the same animal. However, the variability across subjects can be substantial, depending on both the frequency of activity and its overall duration. Similarly, heart rate variability in human subjects is studied by recording multiple cardiac cycles using electrocardiography (ECG) and monitoring changes in the time between cycles (RR intervals) [3]. The number of RR intervals recorded from a single subject can be in the tens to thousands, varying widely depending on both heart rate and the total duration of the experimental protocol.

Such physiological studies share several challenges, including that there are multiple observations 1 recorded from a single subject and that not all subjects have the same number of observations (i.e. an unbalanced design). In some studies, data from subjects in a given group are pooled and the group sample size is taken as the total number of observations. This is problematic for two reasons. First, if there are a different number of observations for each subject, this means that subjects will be weighted differently in the pooled sample. If there are subjects that present different behaviors but have a large number of observations, the pooled sample could skew towards these subjects. The issue becomes more serious if some of the variability between subjects in terms of the number of observations depends on one of the factors under study. For example, in studies of exercise, the number of cardiac cycles in a subject will depend on their heart rate. If observations are pooled, the distribution could skew towards those subjects with higher heart rates. Statistically, the group sample size should be the number of subjects, with each subject represented only once. This brings us to a second problem, namely that many statistical tests used to compare groups assume that the data points are independent of one another. However, multiple observations from the same subject are usually not independent, and treating them as such constitutes pseudoreplication [4].

Though the importance of pseudoreplication continues to be debated [5], it does appear to be prevalent in some disciplines (e.g. 12-36% in neuroscientific studies [6]) and can lead to “erroneous positive results” in physiological studies [7]. Analysis strategies and statistical tests exist to handle repeated measures, but are not ideal for the types of data discussed here. For example, one could take a single measure (e.g. the median RR interval) for each subject, and then work with a distribution of medians for the group – a method known as aggregation [8]. This addresses the pseudoreplication problem, but involves losing a lot of the richness in the data, among other issues [8]. Statistical tests like repeated measures ANOVA are not suitable, since these have assumptions, e.g. about sphericity and completeness of data [9], which are often not met.

To address these issues, we have developed a method designed to preserve the spread of the data from each subject, construct a representation of the group distribution using the distributions from its individuals, and still guarantee that each subject is represented only once in the analysis. Importantly, this method does not require the data to be approximately normally distributed, or that the range for the data points is restricted to any domain. We illustrate this method by using it to analyze two physiological datasets. The first consists of intramuscular recordings taken from *Drosophila* larvae during fictive crawling. These data were obtained by one of the present authors (ECM) and have been described previously [2]. The second dataset consists of ECG recordings from volunteer human subjects taken before, during, and after exercise. These recordings were obtained by two of the present authors (JHAR and ATP) and have not been reported previously. We share all the data and code used to implement these analyses. Our results show the simplicity and broad applicability of this method, and we hope it will provide a viable alternative to pooling for researchers analyzing similar physiological data.

## Methods

### Physiological data

#### *Drosophila* muscle recordings

The fly lines, rearing conditions, dissection method, and electrophysiological recording technique have all been described in detail previously [2]. Briefly, larvae were dissected to expose the body wall muscles, and flexible sharp glass electrodes were inserted into muscle 1 (M1) [10] to record intracellularly from neighboring abdominal segments. In this way, spontaneous bouts of peristaltic muscle activity (fictive crawling) can be recorded as waves of contraction that advance from the posterior to anterior body segments. This activity is registered as regular bursts of action potentials. While several measures can be obtained from these data, we focus on the cycle duration, which is the time elapsed from the start of one burst to the start of the next. We compared the cycle durations of two groups: (1) wildtype (WT) larvae (n=21), and (2) larvae expressing the *slo*^1^ mutation (n=10; Bloomington Drosophila Stock Center No. 4587; RRID BDSC_4587). Again, effects of this mutation on bursting activity have been reported previously in [2].

### Human ECG recordings

Subjects were student volunteers between 18 and 24 years of age recruited within the School of Sciences at the National Autonomous University of Mexico (UNAM). Since this was a non-invasive observational study performed with volunteers, and the School did not have a bioethics review board at the time, no institutional permissions were required. However, all volunteers read and signed a letter of consent to participate in the study. ECGs were recorded using a Zephyr BioHarness before, during, and after exercise on a stationary bicycle (Pro-Form 325 CSX). Subjects were asked to remain at rest for the first 7 minutes of the recording, then begin cycling, gradually increasing intensity during minutes 7-9. At the 9-minute mark, subjects were asked to further increase their exercise to achieve a pre-specified intensity level (e.g. 60 watts, as indicated on the bicycle interface), and then maintain that intensity until minute 20. A recovery period followed until around the 30-minute mark, when the recording ended. The time elapsed between successive cardiac cycles was measured as the RR interval. We compared the RR intervals of two groups: (1) volunteers classified as physically fit, or active (n=11), and (2) volunteers classified as less physically fit, or sedentary (n=10). Classification was based on exercise tolerance: subjects who reached a heart rate of 150 beats per minute (BPM) at the 60-watt exercise intensity level were classified as less fit (group 2) than subjects whose heart rate did not reach 150 BPM until the 75-watt level (group 1). Based on our protocol, subjects did not proceed to the next watt-level once reaching 150 BPM. Therefore, for this study, we compared recordings at the maximum watt-level achieved by all subjects, i.e. 60 watts. To also minimize variability, we compared subjects’ RR intervals during the period of exercise that was most stable, i.e. minutes 17-20. (Changes in RR intervals during the other protocol time periods will be explored in a future paper.)

## Data analysis

Let *X* represent a continuous random variable of interest (e.g. RR interval).

### Empirical distributions

Consider a sample *S* = (*x*_1_, …, *x*_*n*_) and assume, without loss of generality, that *x*_*i*_ ≤ *x*_*i*+1_ for all *i* ∈ {1, …, *n* − 1}. Assuming the random variable *X* is continuous, the empirical distribution function (EDF, [11, 12]) for the sample can be defined as

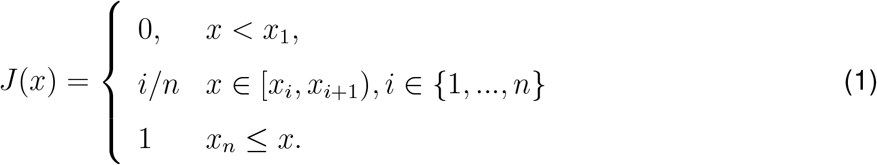

If the sample is unbiased and contains enough data points, the function *J* (*x*) can serve as an approximation for the distribution function of the variable represented by the points in *S* [12].

### Group samples

Assume that two groups of samples *C* = {*C*_1_, …, *C*_*M*_} and *D* = {*D*_1_, …, *D*_*N*_} of *X* values are collected, and let *F*_1_, …, *F*_*M*_ and *G*_1_, …, *G*_*N*_ respectively represent the EDFs from the samples of the two groups. The point-wise averages of the EDFs for the two groups can then be calculated by letting

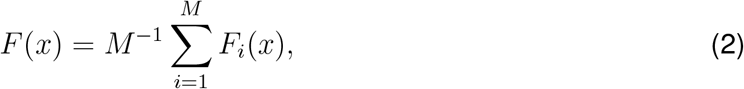

And

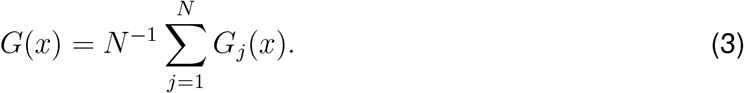

These point-wise average functions *F* and *G* can be thought of as representative EDFs for the groups *C* and *D*, respectively. For the cycle durations (CDs) obtained from *Drosphila* muscle recordings, the sample EDFs from the two groups of subjects will be denoted as *F*_*CD*_ (WT) and *G*_*CD*_ (Slo), respectively. Similarly, for the RR intervals obtained from human subjects, the two groups will be respectively labeled *F*_*RR*_ (active) and *G*_*RR*_ (sedentary).

For the two groups in our case, the whole range for the variable *X* can be defined as

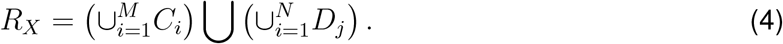

The interval [*a, b*] with *a* = min {*x* ∈ *R*_*X*_} and *b* = max {*x* ∈ *R*_*X*_} serves as an extended range for values of *X* as extracted from the two groups. From there, it is possible to evaluate the average EDF for each group using [*a, b*] as a common interval.

### Sampling

Kolmogorov’s theorem [13] can be used to obtain values representative of a random variable *X*. If *F* is the distribution function of *X*, and *p* ∈ [0, 1], then *q* = *F* ^−1^(*p*) is a value that represents *X* so that *P* (*X* ≤ *q*). Therefore, any ordered sample *α*_1_, …, *α*_*n*_ uniformly drawn from the interval [0, 1] can be used to sample *n* values for *X*. Explicitly, the sample would be *x*_1_ = *F* ^−1^(*α*_1_), …, *x*_*n*_ = *F* ^−1^(*α*_*n*_). For instance, the set {0, 0.25, 0.5, 0.75, 1} yields the group mínimum, maximum, and the quartiles for *X* as described by *F*. This procedure can then be applied using a large enough number of points uniformly sampled from the interval [0, 1] to obtain representative samples of the two groups from the two average EDFs in equations (2)-(3).

### Histograms

To visually explore how the data are distributed and any potential shifts between groups, we construct histograms using samples drawn from each group EDF. For any given number of bins *n*, it is then possible to create a uniform partition *a* = *p*_0_, …, *p*_*n*_ = *b* of the interval of [*a, b*], with each subinterval having length (*b* − *a*)*n*^−1^. The size of the subintervals can be selected according to physiological or experimental criteria. Histograms are only used herein for illustrative purposes and not for any statistical testing. As such, the chosen bin size will not affect the results.

### Statistical testing and sensitivity analysis

Group distributions can be overlayed to visually demonstrate shifts in the observed values as a result of certain group characteristics or treatments. However, statistical testing is needed to determine whether these shifts are significant relative to a predetermined margin of error. We elected to use the Mann-Whitney U test (also known as the rank sum test) due to its lack of assumptions about the distribution of the data and its ability to test for “distributional differences”, including changes in shape and spread [14]. The alpha value for the test was set at 0.05. We test the null hypothesis that the *X*-values from the two groups come from the same distribution.

The values entered into the rank sum test were calculated from the group EDFs using the sampling method described above, i.e. by evaluating the EDFs for each of the groups at *n* uniformly chosen values between 0 and 1. To determine the effect of selecting a specific *n*, we performed a sensitivity analysis. First, two groups were selected for comparison (*Drosophila* WT vs Slo, or human active vs. sedentary). Then, we evaluated the EDFs for each group using an *n* that corresponded to theoretical sample sizes of between 5 and 40 values with the aid of a random number generator. Finally, a rank sum test was performed to compare the two reconstructed samples, and the resulting p-value recorded. This process was repeated ten times for each theoretical sample size, so that p-values could be compared to determine the consistency of the results.

### Analysis software

Burst times were extracted manually from *Drosophila* muscle recordings using Spike2 software and exported to csv files, as explained in [2]. RR intervals were extracted from human ECG recordings using the Zephyr BioHarness software (v.1.0.24.0) and exported to txt files. Code to analyze csv or txt data was written in Python 3.13.3 using functions from NumPy [15] and SciPy [16]. Figures were produced with matplotlib [17] and seaborn [18]. Additional Python modules used include csv, math, os, pyabf, pylab, stats, and sys.

### Resource availability

Resources generated by this study (data, code, figures, etc.) are available via GitHub https://github.com/emckiernan/physio-analysis and archived with a DOI via Zenodo https://doi.org/10.5281/zenodo.1574085. To facilitate reuse, resources are shared under open licenses (see LICENSE.md in our repo). To promote reproducibility, Python code is embedded in Jupyter notebooks [19] that explain the code, how to use it, and how to generate figures.

## Results and Discussion

### Similarities between the datasets

Visual exploration shows that the two datasets share several characteristics important for their analysis (Fig. 1). First, both are continuous time series representing rhythmic physiological processes where the measure of interest repeats at regular intervals (Fig. 1A,B). Second, the measure of interest calculated from each time series (CD or RR interval) captures the length of a single cycle (Fig. 1C,D). Third, because we are measuring cycle length and the cycle repeats, the total number of observations from a single subject in both datasets will depend on two factors: (1) the frequency of activity (i.e. how many cycles occur per unit time), and (2) the total time of recording. Even when we can control the second factor by setting a fixed recording time, we will still have different numbers of observations across subjects due to variability in the first factor. Finally, because of these factors, subjects may have widely varying numbers of observations, with subjects generating higher-frequency activity necessarily having larger numbers.

**Figure 1:**
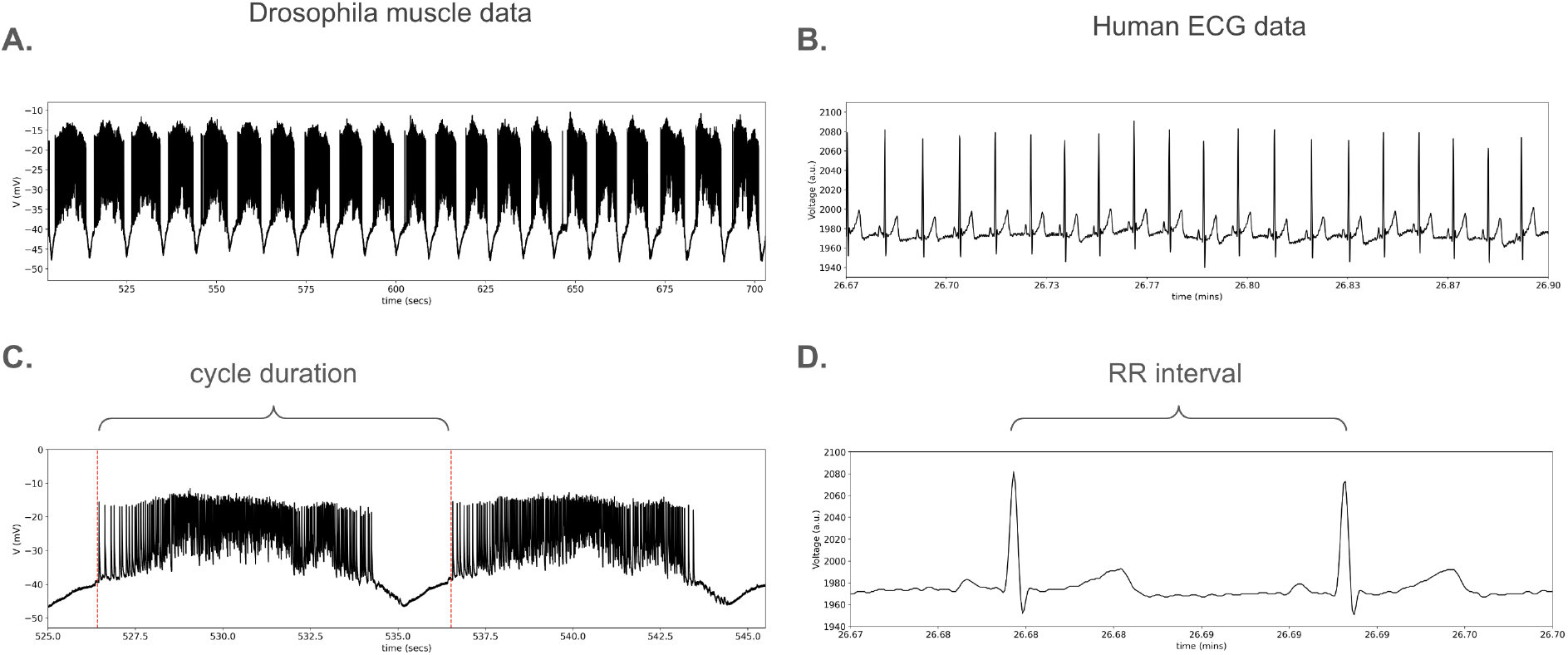
Physiological data. (A.) Intracellular muscle recording from a WT *Drosophila* larva. Activity was recorded during posterior to anterior contractile waves in muscle 1. Data from ECM [2]. (B.) ECG recording from a human volunteer. Original data from JHAR and ATP. (C.) Zoom-in of data in A showing how cycle duration is calculated. (D.) Zoom-in of data in B showing how RR interval is calculated.

### Problems with pooling

Also in both datasets, we end up with repeated observations from a single subject that are not independent of one another. Many studies of such physiological processes ignore this issue, and pool observations for all subjects belonging to a given group (e.g. all CDs for WT subjects, or all RRs for active subjects). To understand why this could be problematic, we look at the distributions for each subject in our samples to see their relative contributions. We do this by plotting their kernel density estimate (KDE), which is essentially a smoothed version of the histogram [20] for each subject that makes comparison visually easier (Fig. 2A,B). For both datasets, the subject KDEs show variability in their amplitude, shape, and spread. In other words, individual subjects may contribute very different sets of values to a pooled distribution. For example, the data from the WT *Drosophila* larvae show that two subjects have larger and more left-shifted KDEs than the rest of the subjects in the sample (see arrows in Fig. 2A). These are the two subjects with the largest numbers of observations (49 and 62 bursts). If we pool the data, these two subjects will be over-represented and could pull the group distribution to the left. We see something similar in the RR data from the active group (Fig. 2B). There is one subject whose values fall far to the right of the other subjects (see arrow), and could pull the distribution in that direction.

**Figure 2:**
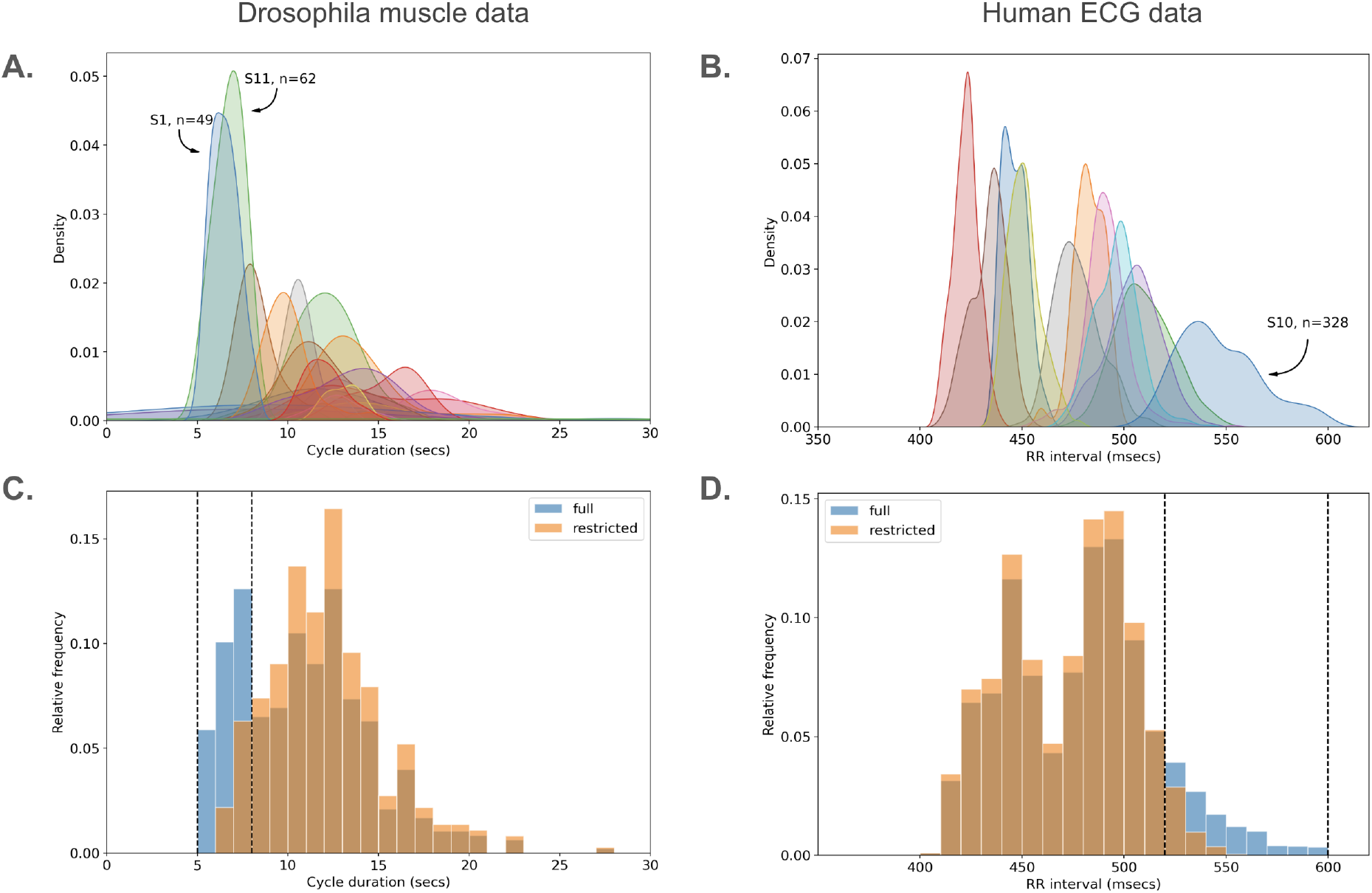
Effects of pooling data. (A.) KDEs for each subject in the WT *Drosophila* sample. Arrows mark two subjects with left-shifted peaks. (B.) KDEs for each subject in the active volunteer group. Arrow marks the right-shifted subject. (C.) Histograms of *Drosophila* data in A, comparing the full pooled sample (blue) to the restricted sample with two subjects removed (orange). (D.) Histograms of the active volunteer data in B, comparing the full pooled sample (blue) to the restricted sample with one subject removed (orange). Darker brownish areas in C and D show overlap between the blue and orange bars. Dashed lines in C and D delineate areas of difference between the full and restricted histograms.

How much could these subjects affect a pooled distribution? To determine this, we took the same groups (WT *Drosophila* or active human volunteers), pooled subjects’ observations within those groups, and plotted the respective histograms (referred to as the ‘full’ samples). We then removed the subjects of interest (those marked with arrows in Fig. 2A,B), and compared the histograms with these subjects removed (‘restricted’ samples). The results are similar for the two datasets (Fig. 2C,D). For the WT *Drosphila* larvae, removing the two subjects caused the three left-most bars in the histogram to either disappear or greatly diminish. In fact, this removed an entire peak which made the distrbution (at least at this bin width) look bimodal. For the active volunteers, removing the one subject eliminated or reduced a large chunk (8 bars at this bin width) of the right side of the histogram. These are well-known effects of what might be considered outliers, but we include them as visual confirmation of how individual (or few) subjects can shift a pooled distribution, especially when samples are small.

### Technique based on EDFs

Having established that pooling observations at best violates the assumption of independence and at worst leads to skewed distributions, we need a better way to deal with such data. As described in the Methods, we did this by first calculating the empirical distribution function (EDF) of the CD or the RR values for a single subject, and then calculating the point-wise average over all subjects in a given group to calculate the group EDF (Fig. 3). In this way, we do not lose the richness of the data – each subject is represented by their EDF, which captures the full range of values in their observations, rather than a single value like a mean or median. At the same time, taking a point-wise average of the EDFs means that at each point, each subject is represented only once. The subject EDFs are all independent of one another, and therefore, the resulting averaged group EDF is made up of independent ‘observations’ (subject EDFs), thereby avoiding pseudoreplication. In other words, each subject now contributes equally to the group, regardless of their initial number of observations. Importantly, while outliers could still skew a group distribution, using this averaged approach means no one subject is more able than any other to pull the distribution solely based on their number of observations. In both datasets, we can see that there is a clear separation between the group EDFs (Fig. 3) that we should be able to test for statistical significance.

**Figure 3:**
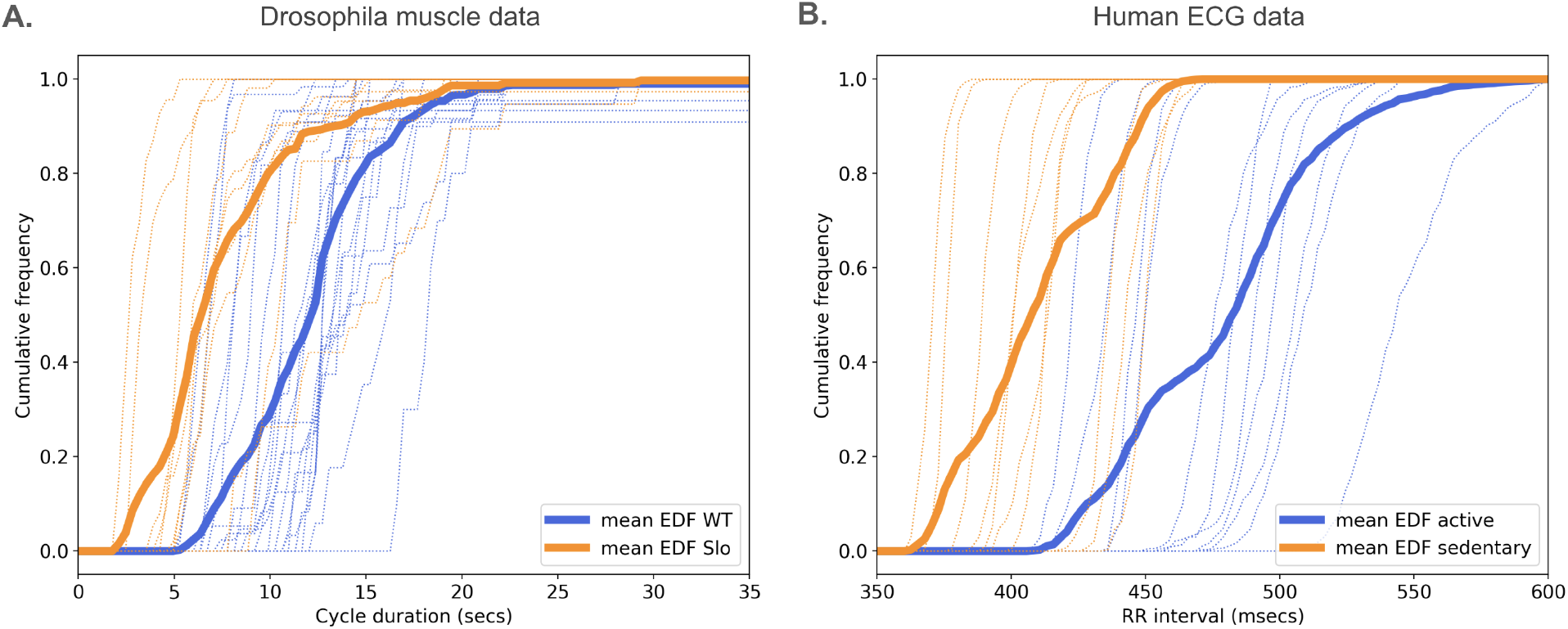
Single-subject and group EDFs. A. Single-subject (thin dashed lines) and averaged group (thick solid lines) EDFs for WT (blue) and Slo (orange) larvae. B. Single-subject (thin dashed lines) and averaged group (thick solid lines) EDFs for active (blue) and sedentary (orange) human volunteers.

To test for statistical significance, we used the Mann-Whitney or rank sum test with a reference value for significance of 0.05 (standard for physiological studies). As described in the Methods, subsections ‘Samples’ and ‘Statistical testing and sensitivity analysis’, our technique relies on extracting values from the group EDFs for testing. We therefore ran a sensitivity analysis to see how the number of extracted values affects the results. To do this, we first set the theoretical sample size (the number of extracted values) between 5 and 40 values, and then used a random number generator to extract that number of values at different points along the averaged group EDFs. We run the statistical test on these extracted samples, and repeat the process 10 times for each sample size. Graphing the results shows how many of these test-runs hit below the established alpha value of 0.05 for a given sample size (Fig. 4). For the *Drosophila* data, there is a large fluctuation in p-values at the lowest theoretical sample sizes of 5-10 (Fig. 4A). Between 10 and 20, there is still some fluctuation, though steadily decreasing as the sanple size increases. Finally, most if not all test runs are below 0.05 as the sample size goes above 20. Similarly, for the human ECG data, there is some variability in p-values when the sample size is betwee 5 and 10, though less than for the *Drosophila* data, Above a sample size of 10, most if not all runs are well below 0.05 (Fig. 4B). All our real samples include at least 10 and as many as 21 subjects in each group. Therefore, extracting 10-20 values from the group EDFs for statistical testing appears to be both reasonable and conservative. However, the reasonable range of values to extract will depend on the dataset, and should be evaluated on a case-by-case basis.

**Figure 4:**
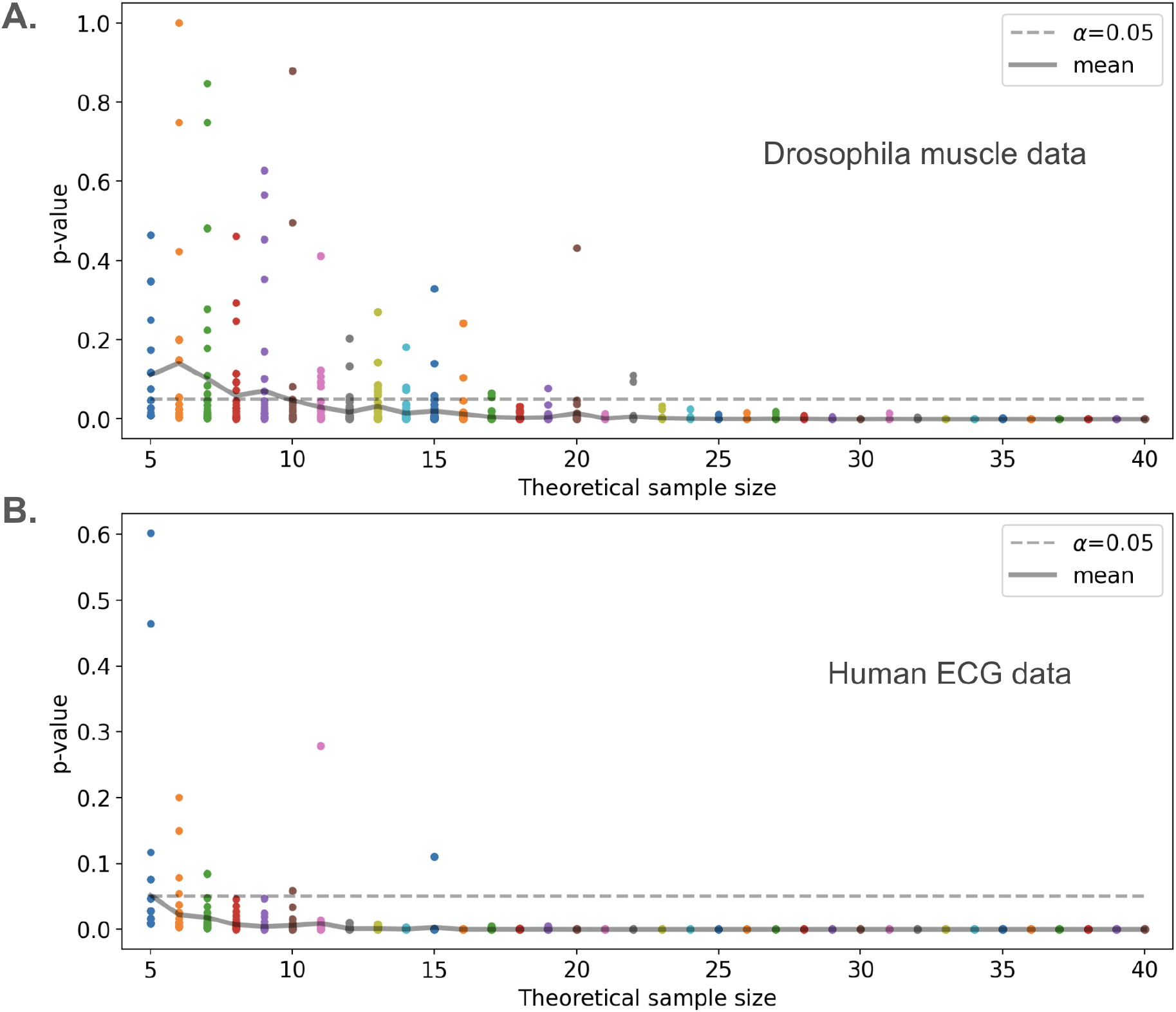
Sensitivity analysis. Statistical testing run 10 times for each theoretical sample size of 5 to 40 values extracted at random points along the group EDFs for *Drosophila* (A.) and human (B.) data. Note that not all points corresponding to each run are visible for all sample sizes due to overlap. Dashed line marks a cutoff for significance at 0.05. Solid line indicates the mean p-value over 10 runs at a given sample size.

## Conclusions

Proper data analysis, including ensuring we meet the assumptions of any statistical tests used, is crucial for the validity of study results and conclusions. We present a technique that allows researchers to analyze physiological datasets which represent positive definite random variables that are not approximately normally distributed, and include uneven numbers of repeated measures across subjects. Our technique has advantages over pooling data, including maintaining independence of measurements used for testing and avoiding pseudoreplication. It also represents advantages over techniques like aggregation in that the original richness of the data is preserved and considered. We also show that this technique can be applied to different datasets, ranging from insect muscle to human heart recordings, demonstrating its generalizability. All our analysis code is publicly available so that it can be reused by other researchers and modified to fit their needs.

